# Single subunit degradation of WIZ, a lenalidomide- and pomalidomide-dependent substrate of E3 ubiquitin ligase CRL4^CRBN^

**DOI:** 10.1101/595389

**Authors:** Helen H. Yu, Justin M. Reitsma, Mike J. Sweredoski, Annie Moradian, Sonja Hess, Raymond J. Deshaies

**Author notes:** AbbVie Clinical Pharmacology Unit, 480 US-45, Grayslake IL 60030. AstraZeneca- MedImmune, LLC, One MedImmune Way Gaithersburg, MD 20878.

## Abstract

Immunomodulators (IMiDs) are an effective class of drugs used to treat blood cancers. IMiDs are believed to work by recruiting protein targets containing a β-hairpin motif for ubiquitination by E3 ubiquitin ligase complexes composed of cereblon (CRBN), Cullin-4a (CUL4a), DNA Damage Binding protein-1 (DDB1), and Ring Box-1 (RBX1). The ubiquitinated protein is subsequently degraded by the proteasome. By characterizing the repertoire of proteins that show an increased physical association with CRBN after IMiD treatment, we identified a novel IMiD substrate, Widely Interspaced Zinc Finger Motifs (WIZ). WIZ contains a C2H2 zinc finger domain, like several other substrates that were previously characterized. We demonstrate that IMiDs stabilize physical association of WIZ with CRBN, deplete WIZ steady state protein levels in a way that is dependent on E3 ligase activity, and enhance the rate of its degradation. Notably, proteins that assemble with WIZ are co-recruited to CRBN by IMiDs but are not degraded, illustrating the potential of targeted protein degradation to eliminate individual subunits of a protein complex. These findings suggest that systematic characterization of the full repertoire of proteins that are targeted for degradation by IMiD compounds will be required to better understand their biological effects.

**Synopsis:** Proteolysis Targeting Chimeras (PROTACs) can be used to precisely target a subunit of a transcriptional complex for degradation in subpopulations of cells.

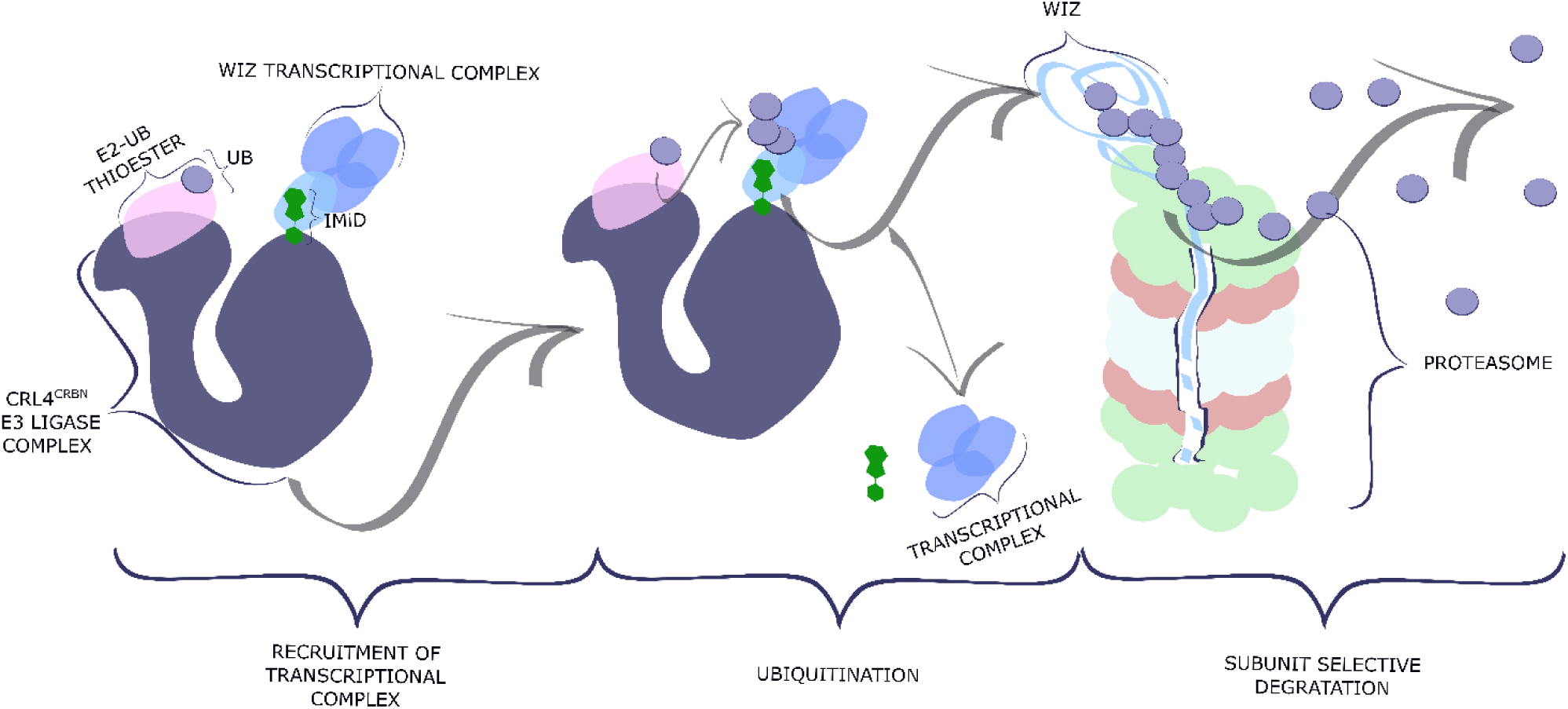

## Introduction

Redirecting the existing cellular degradation machinery to remove unwanted components rapidly and specifically is potentially a versatile therapeutic approach (1). One possible strategy to achieve this goal is to redirect the proteasome, a large enzyme that digests proteins. Selective elimination of one subunit of a multi-protein complex by this approach is possible because the proteasome can extract a single subunit from a complex, unfold it, and thread it into its inner chamber, where the protease active sites reside (2). Therefore, what is needed is a mechanism to redirect the proteasome to remove a therapeutic target in addition to its endogenous substrates.

The proteasome normally identifies and enriches for proteins that need to be removed by using ubiquitin as an affinity tag (3, 4). Ubiquitin is an 8 kDa protein that is ubiquitously expressed in eukaryotic organisms (5). The C-terminal glycine on ubiquitin is conjugated to the amine on lysines of the target proteins via an isopeptide linkage. Formation of an ubiquitin polymer with at least 4 monomeric units yields a signal that binds tightly to the proteasome (6, 7). Therefore, one way to degrade an intracellular therapeutic target would be to catalyze the ubiquitination of the target to redirect it to the proteasome.

Three enzymes act in tandem to catalyze selective ubiquitination of proteins (8, 9). E1 first converts the C-terminal carboxylic acid on ubiquitin into a more reactive thioester to enhance the rate of reaction, as the C-terminus of ubiquitin is fairly chemically inert. The activated ubiquitin is transferred as a thioester to an E2, and the E2 ubiquitin thioester then binds to an E3 enzyme, which also binds to substrate. RING-type E3 enzymes position the E2 for subsequent transfer of ubiquitin to a lysine residue on the bound substrate (10). Since E3 ubiquitin ligases are the components of this pathway that identify which proteins get ubiquitinated, one strategy to redirect the proteasome to digest a therapeutic target is to modify the substrate specificity of an E3 ubiquitin ligase to recognize the protein of interest.

One class of E3 ubiquitin ligases that have been successfully redirected towards a therapeutic target (i.e. neosubstrate) are cullin-RING type E3 ubiquitin ligases (CRLs) (11-21). A distinguishing feature of CRLs is that they contain interchangeable subunits that recruit specific substrates to the enzymatic core (10). CRLs work by catalyzing the discharge of ubiquitin from E2 onto a lysine residue of a juxtaposed natural or neosubstrate.

Investigating molecules that work by re-directing CRL activity toward neosubstrates can inform the design of future drugs. One class of drugs that redirect CRL activity towards neosubstrates is referred to as IMiDs, or immunomodulatory drugs (24). IMiDs have been shown to be effective for the treatment of hematological malignancies by redirecting cereblon (CRBN), the interchangeable substrate receptor that determines the specificity of the CRL4^CRBN^ complex, to recruit a set of neosubstrates (Fig.1a) (14, 20, 21, 25). CRBN is one of many substrate receptors that are recruited to CUL4-RBX1 by DDB1 to form CRL4 complexes (Fig.1a) (22, 23).

**Fig. 1.**
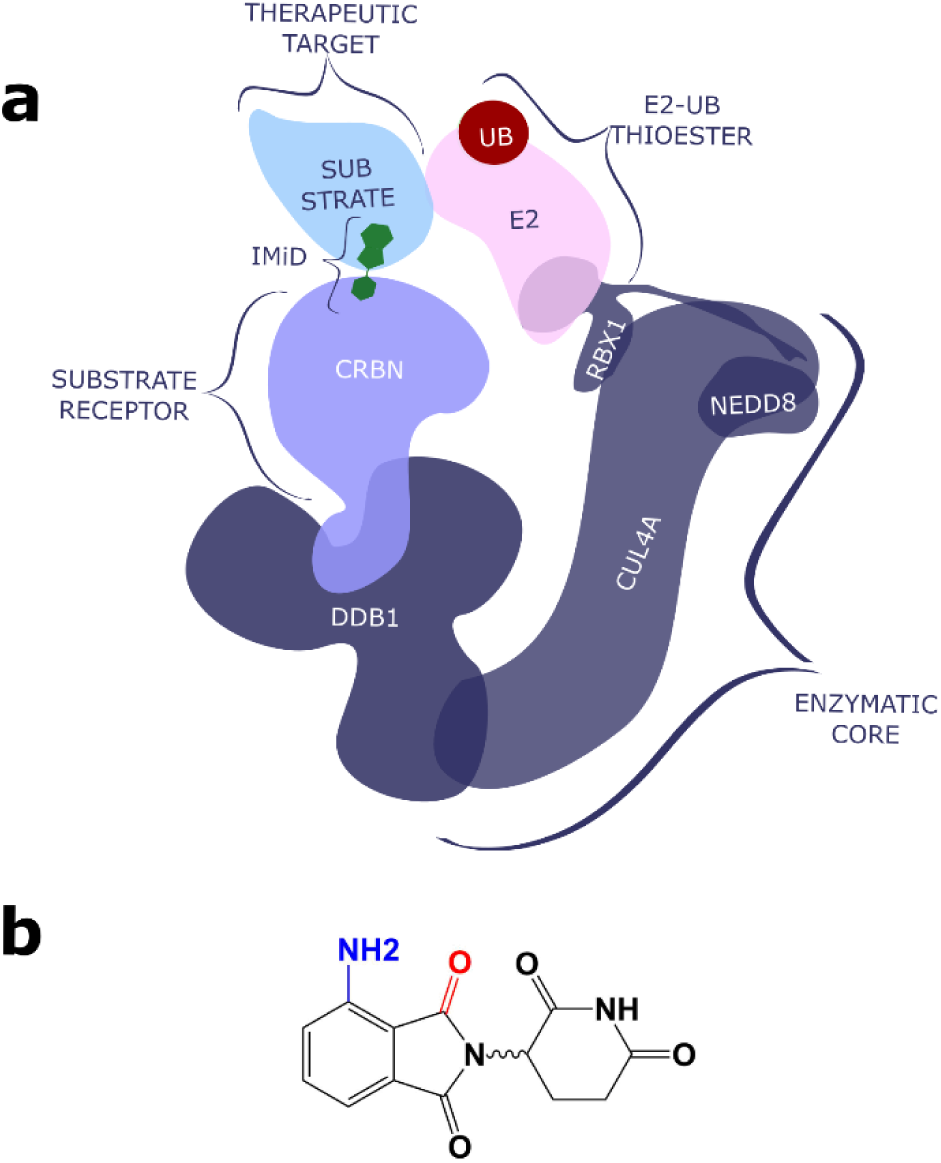
Mechanism of action for IMiDs. (**a**) CRBN is normally in complex with DDB1, which in turn interacts with CUL4-RBX1 to form the ubiquitin ligase CRL4^CRBN^(22, 23). IMiDs nucleate a novel protein-protein interaction between CRBN and a cellular protein (neo-substrate) (14, 20, 21). Ubiquitin-thioesterified to an ubiquitin conjugating enzyme (E2) is recruited to CRL4^CRBN^ by RBX1, which brings activated ubiquitin into proximity of the neosubstrate (10). (**b**) General structure of IMiDs. Addition of the blue NH_2_ group to thalidomide yields lenalidomide. Removal of the red carbonyl group from lenalidomide yields pomalidomide.

**Fig. 2.**
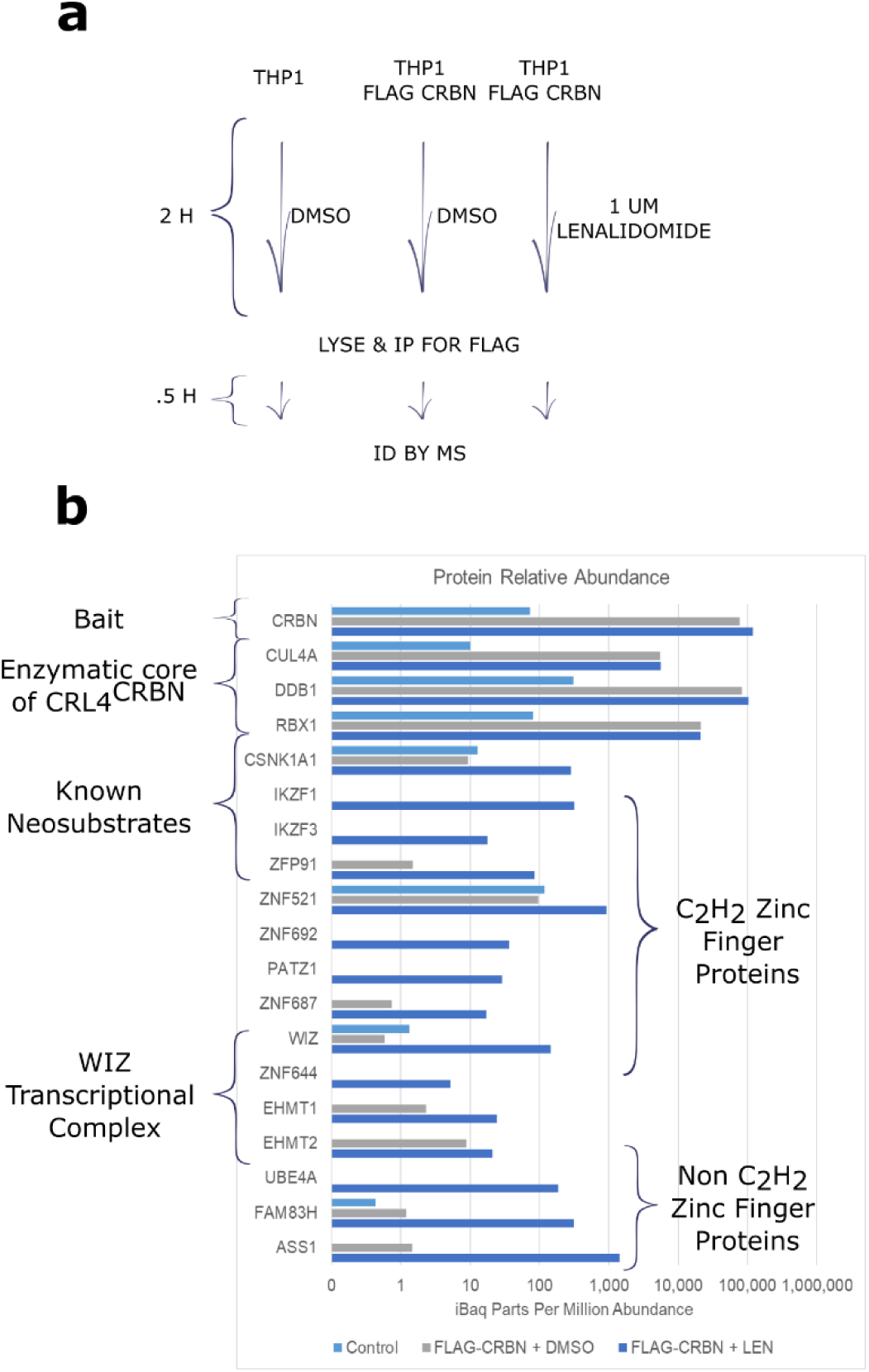
Proteins with Zinc Finger Domains show enhanced physical association with CRBN in the presence of lenalidomide. (**a**) Outline of IP-MS strategy to identify proteins that show an enhanced physical association. (**b**) Peptide count of proteins identified in the experiment. This includes those known to bind CRBN, known core components of the CRL4^CRBN^ complex, zinc finger proteins, and novel binding partners.

IMiDs are bipartate molecules that act as a molecular bridge to link neosubstrates to CRBN. The glutarimide moiety binds CRBN, and the phthalimide moiety binds to protein targets (Fig.1b) (26, 27). All known IMiD-dependent neosubstrates reported for CRL4^CRBN^ are predicted to contain a β-hairpin motif, like Casein Kinase 1*α* (CK1*α*) and the C_2_H_2_ zinc finger proteins Ikaros (IKZF1), ZFP91, Sal-like protein 4 (SALL4) and Aiolos (IKZF3) (Fig.1a) (14, 20, 21, 25, 28, 29). Crystal structures have shown that the β-hairpin motif comes into contact with the phthalimide moiety of IMiDs bound to CRBN, which explains how these drugs work (30, 31).

Because novel substrates are recruited to CRBN through the phthalimide moiety of IMiDs, understanding how modification of the basic bicyclic IMiD scaffold influences the identity of the neosubstrates that are recruited to CRBN will be critical for the development of new analogs with different patterns of selectivity (Fig.1b). Interestingly, relatively modest alterations to the IMiD scaffold can have surprisingly dramatic effects. For example, substitution of an amine at the 4’ position of thalidomide (Fig.1b) (as occurs in lenalidomide and pomalidomide) results in a 100 to 1,000-fold increase in potency (32). Another dramatic example of specificity is seen with deletion of a simple carbonyl from pomalidomide to generate lenalidomide (Fig.1b), which enables targeting of the neosubstrate CK1α for ubiquitination and degradation (25, 30). Thus, relatively subtle modifications to the IMiD chemical scaffold result in dramatic changes in its biological properties. Additionally, it remains unknown how the effects of IMiDs are modified in different cell types. If we are to understand how to harness targeted protein degradation, it will be important to understand the physical and chemical basis for targeting by identifying and characterizing specific neosubstrates.

## Results

### IMiDs enhance physical association of CRBN with proteins with Zinc Finger domains

Previous IMiD substrates have been identified in proteomics experiments by determining which proteins accumulate as ubiquitinated intermediates or exhibit a shorter half-life in the presence of IMiDs (19, 21, 25, 28, 29). However, there are several disadvantages to both of these methods. First, ubiquitinated intermediates typically account for a tiny fraction of the total protein, and only a tiny fraction of all peptides from a protein have ubiquitin conjugated to them (33). Second, the rate of decay is nonlinear, and the half-life of proteins can range from minutes to months (34), potentially confounding a large-scale screen analyzed at a single time point.

To circumvent these issues, we chose to identify proteins whose physical association with CRBN was enhanced upon treatment with lenalidomide, as we reasoned that this was the most direct and sensitive method. To accomplish this, we used a THP-1 cell line that was transiently transfected to express FLAG-CRBN (Fig.2a).

We identified candidate proteins by treating cells with or without lenalidomide for 2 hours, immunoprecipitating for CRBN via the FLAG epitope, and identifying which proteins showed enhanced physical association with CRBN in the presence of lenalidomide via quantitative shotgun mass spectrometry (Fig.2a). Components of the enzymatic core of CRL4^CRBN^ (CUL4a and RBX1) and the adapter (DDB1) that scaffolds CRBN’s interaction with the enzymatic core were recovered in equal amounts regardless of the presence or absence of lenalidomide (Fig.2b).

Previously characterized neosubstrates (ZFP91, IKZF1, IKZF3, and CK1*α*) showed IMiD-dependent association, validating this approach (Fig.2b) (14, 20, 21, 25, 28). However, one limitation of this approach is that enhanced physical association with CRBN might not necessarily result in a productive degradation event (35).

### Levels of transcriptional complex member WIZ are reduced upon IMiD treatment

To investigate how many of the IMiD-dependent binding events translated into productive degradation, we examined by immunoblotting a subset of the identified proteins to see if their steady state levels dropped after IMiD treatment (Fig.3a–c). We selected proteins with a zinc finger domain, because multiple previously characterized IMiD-dependent substrates (e.g. IKZF1, IKZF3, ZFP91, SALL4, ZFP692) are zinc finger proteins (14, 20, 21, 25, 28, 29). L363 cells were treated with pomalidomide, lenalidomide, or thalidomide for 12 hours at multiple doses (0, 1, and 10 µM), in biological triplicate (Fig.3a–c). Both isoforms of WIZ (the identity of the isoforms was confirmed by knockdown; Fig. S1) showed the largest fold change after treatment with pomalidomide (Fig.3a). Previously characterized substrates (e.g. IKZF1) showed IMiD dependent depletion (Fig.3a–c). Additionally, CK1*α* exhibited lenalidomide selective depletion, (Fig.3b) as had been reported previously (25).

**Fig. 3.**
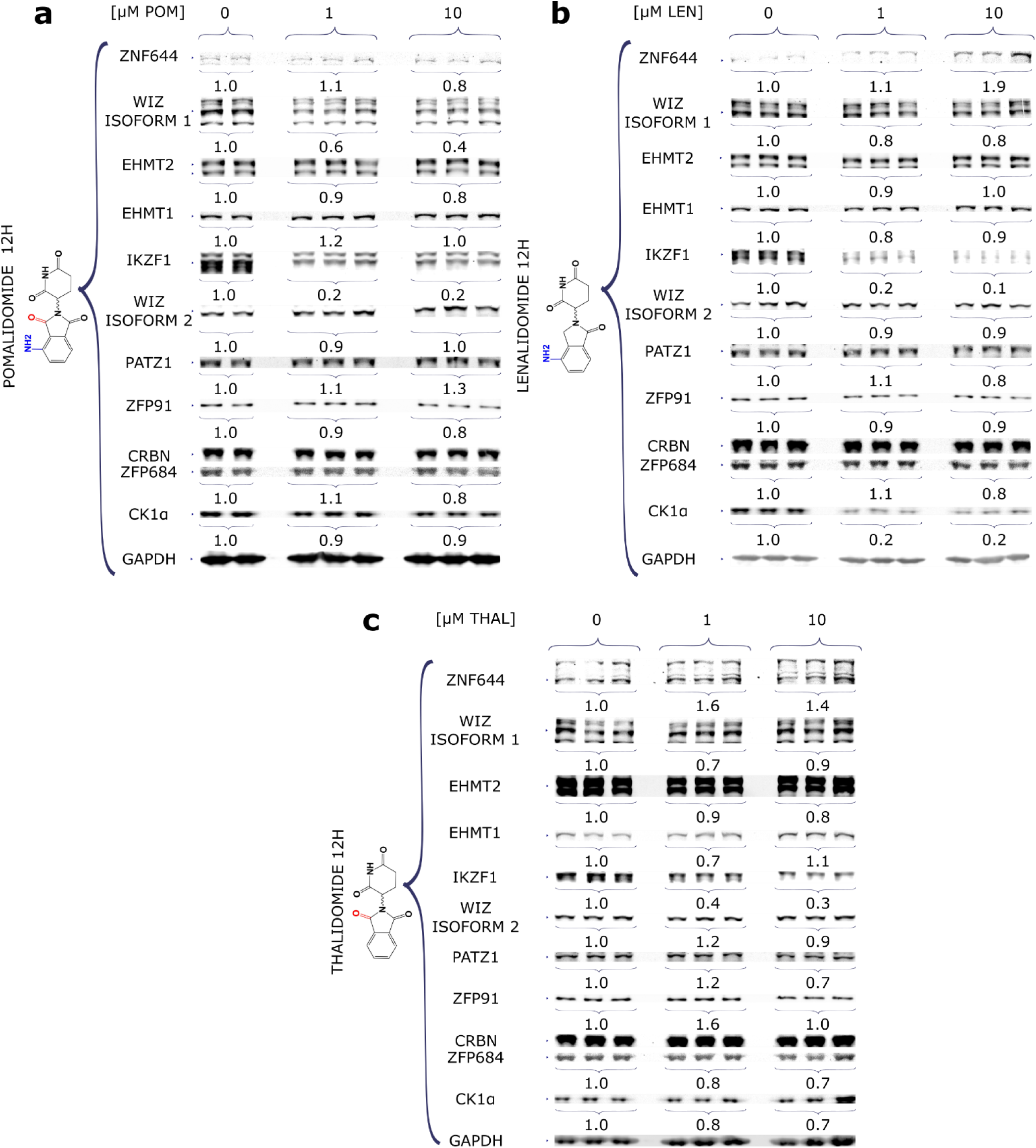
WIZ protein abundance shows IMiD dependent regulation. L363 cells were maintained in RPMI-1640 with 10% FBS and 2 mM glutamine before treatment with (**a**) pomalidomide, (**b**) lenalidomide, or (**c**) thalidomide for 12 hours at 0, 1, or 10 µM. Cell lysates prepared from triplicate cultures were processed for SDS-PAGE and immunoblotting with antibodies to the indicated proteins and the signals were quantified on a LICOR Odyssey. The median level of each protein normalized to no drug is shown below the immunoblot image for each set of triplicates. All protein levels were normalized to GAPDH, which served as a loading control. The previously identified substrates ZFP91, IKZF1, and CK1*α* serve as positive controls (14, 20, 21, 25, 28). PATZ1, ZFP684, and ZNF644 were the other zinc finger proteins screened.

### IMiD regulation shows cell type dependence

To understand how generalizable IMiD-dependent depletion of WIZ was across different cell types, THP-1 cells were treated with pomalidomide and lenalidomide (0, 1, and 10 µM) for 12, 24, 36, and 48 hours and protein levels were evaluated by immunoblotting (Fig.4a–b). Previously characterized IMiD dependent substrates like IKZF1 (14, 20, 25) served as a positive control. Unlike what we observed in L363 cells (Fig.3a), no reduction of WIZ was observed in THP-1 cells (Fig.4a-b), whereas IKZF1 was depleted in both cell types (Figs.4a-b and 3a-c). Previous work had indicated that WIZ was downregulated in H9 hESC (29). Thus, the IMiD effect on WIZ shows cell type dependence.

**Fig. 4.**
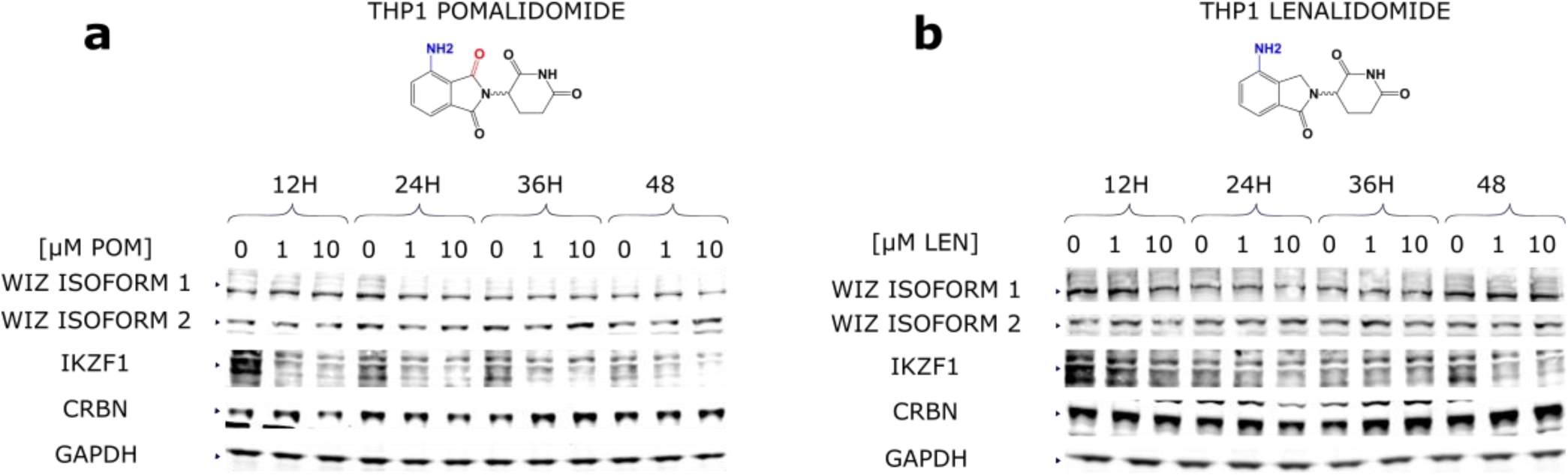
IMiD regulation of WIZ is cell type specific. THP-1 Cells were then treated with (**a**) pomalidomide or (**b**) lenalidomide at 0, 1, or 10 µM. for 12, 24, 36, and 48 hours. Cell lysates were separated by SDS PAGE, and immunoblotted against antibodies for WIZ, CRBN, IKZF1, CK1*α*, and GAPDH.

### WIZ binding partners are not regulated by IMiD

WIZ is a protein that is normally in a transcriptional complex with EHMT1, EHMT2, and ZNF644 (36, 37). It is believed to target methyltransferases to specific DNA sequences via its zinc finger motif. Our mass spec data revealed the entire complex was recruited to CRBN after treatment with IMiD (Fig.2b). To tease out if WIZ was the only component of the transcriptional complex that showed regulation, MM.1s cells were treated with lenalidomide or pomalidomide (0, 1, or 10 µM) for 24 hours, and WIZ, EHMT1, EHMT2, and ZNF644 were evaluated by immunoblotting (Fig.5a–b). Depletion of both WIZ and the validated substrate IKZF1 were observed, but the other components of the WIZ transcriptional complex remained stable.

**Fig. 5.**
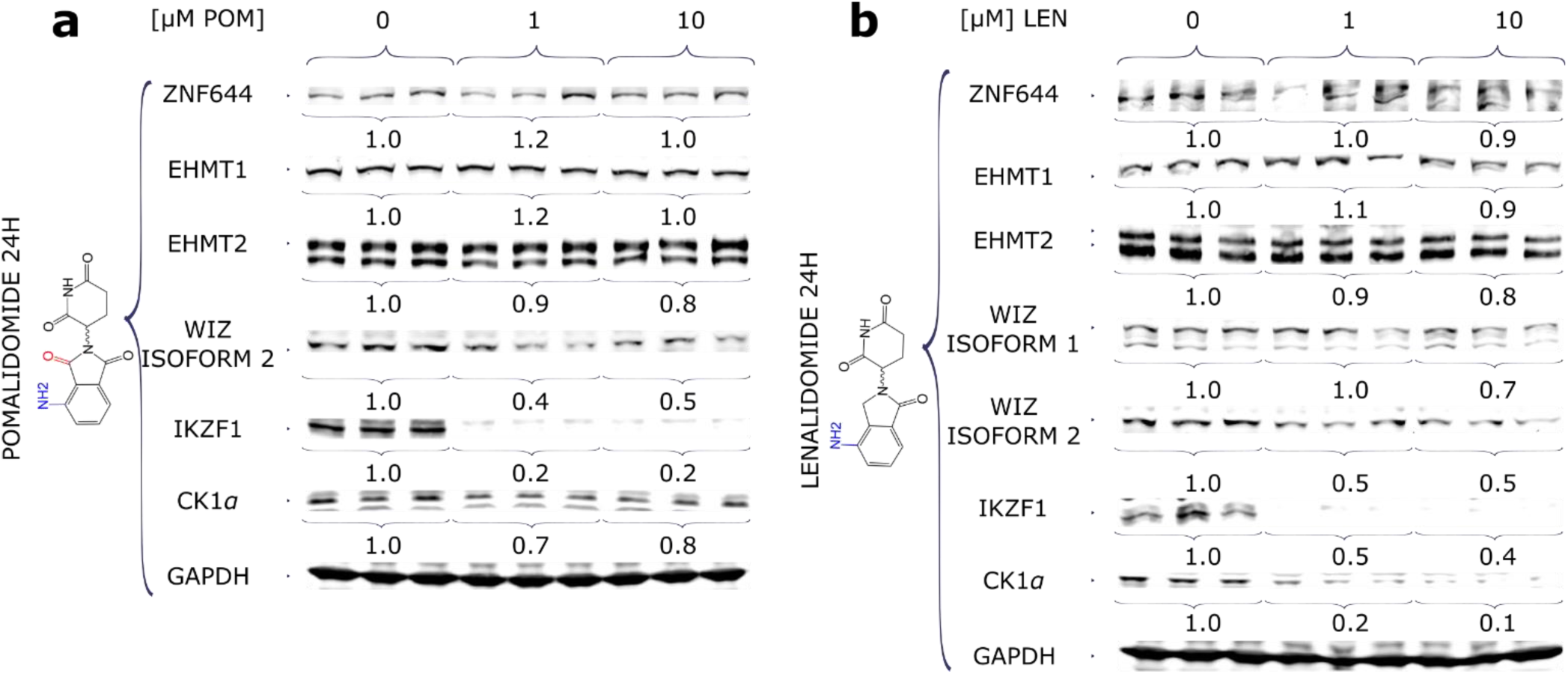
IMiDs regulate the abundance of WIZ, but not the other components of the complex. MM.1s cells were treated in triplicate for 24 hours with (**a**) pomalidomide or (**b**) lenalidomide at 0, 1, or 10 µM. Cell lysates were separated by SDS PAGE and immunoblotted with antibodies against WIZ, EHMT1, EHMT2, and ZNF644. Immunoblot signals were quantified on a LICOR Odyssey. The mean level of each protein normalized to no drug and to the GAPDH signal in the same samples is shown below the immunoblot image for each set of triplicates.

### Levels of WIZ are specifically modulated by IMiDs with an amine modification at the 4’ position

To elucidate how subtle modifications in the IMiD scaffold influenced WIZ depletion, MM.1S cells were treated with pomalidomide, lenalidomide, or thalidomide (0, 1, and 10 µM) for 12, 24, 36, and 48 hours (Fig.6a–c). Both isoforms of WIZ showed the largest fold change upon treatment with IMiDs with an amine in the 4’ position (lenalidomide and pomalidomide), with pomalidomide showing the strongest effect (Fig.6a-b).

### IMiD dependent depletion of WIZ is dependent on CRL4^CRBN^’s E3 ubiquitin ligase activity

IMiDs enhanced the physical association between WIZ and CRBN, resulting in the depletion of WIZ which is consistent with the hypothesis that WIZ is a new CRL4^CRBN^ neosubstrate. We next sought to directly test this hypothesis. Depletion of CRBN in MM.1s cells strongly blocked pomalidomide-induced depletion of WIZ (Fig. 6b, d), confirming that CRBN was indeed required for the effect. To address the role of CRL E3 activity and the proteasome, L363 cells were treated with 0, 1, or 10 µM of pomalidomide for 12 hours in the absence or presence of the Nedd8 conjugation inhibitor pevonedistat or the proteasome inhibitor bortezomib (Fig.7a–c) (38, 39). IMiD depletion of WIZ and known substrates such as IKZF1 was blocked upon cotreatment with proteasome inhibitor, indicating the effect was dependent on proteasome activity (Fig.7a, b). Nedd8 is an ubiquitin-like protein that activates CRLs upon its conjugation to the cullin subunit (41). Co-treatment with 250 nM pevonedistat caused accumulation of CUL4 in the de-neddylated state, indicating that the enzymatic core of the complex was catalytically inactive (Fig.7a, c). IMiD depletion of WIZ and IKZF1 was also blunted by pevonedistat, indicating that the effect was dependent upon the core complex being enzymatically active (Fig.7a, c).

**Fig. 6.**
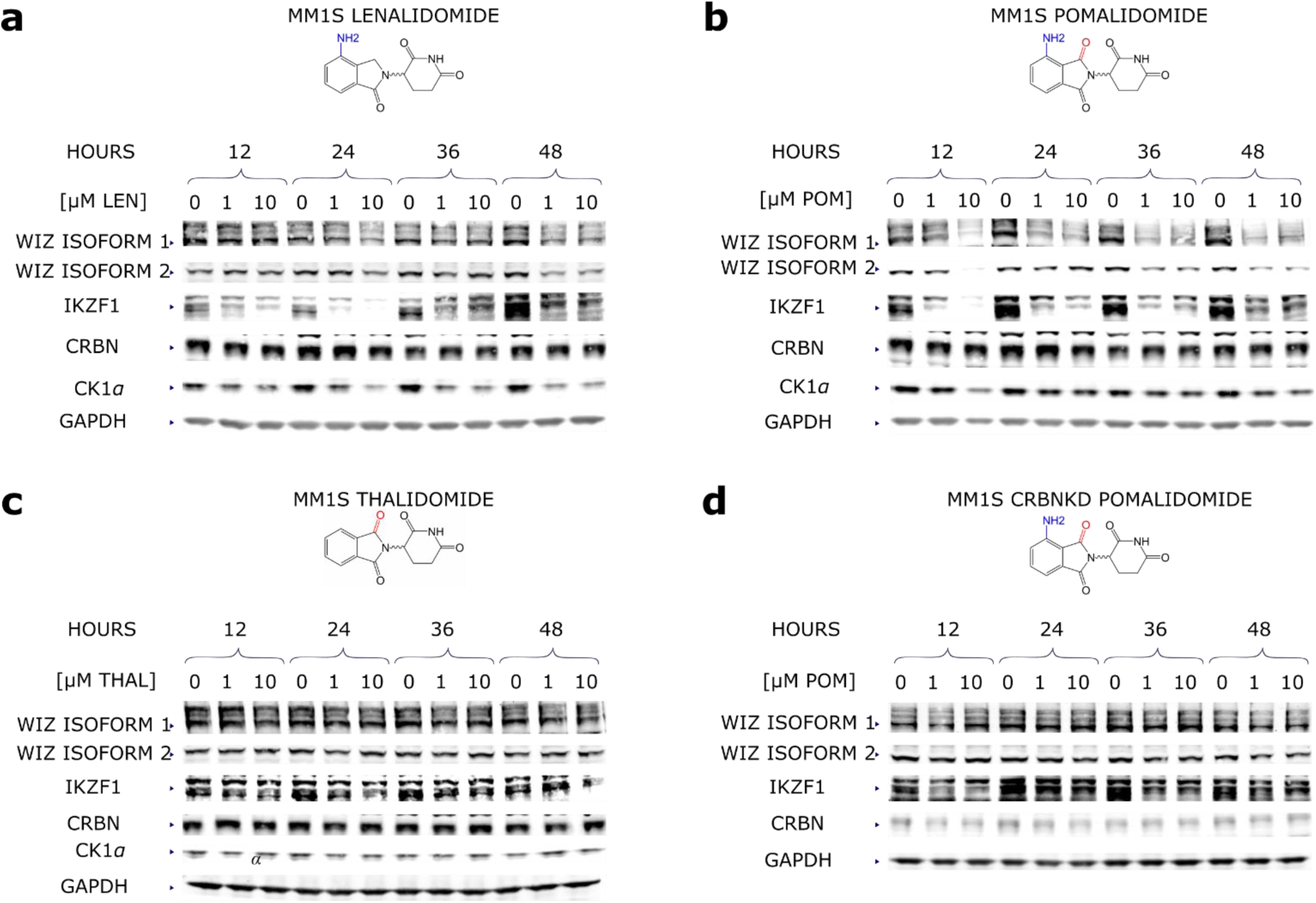
WIZ protein abundance is specific to IMiDs modified with an amine at the 4’ position. (**a**-**c**) MM.1s or (**d**) MM.1s cells with CRBN knocked down with an shRNA were treated with (**a**) lenalidomide, (**b**) (**d**) pomalidomide, or (**c**) thalidomide at 0, 1, or 10 µM for 12-48 hours. Cell lysates were separated by SDS PAGE and immunoblotted against antibodies for WIZ, CRBN, IKZF1, CK1*α*, and GAPDH.

**Fig. 7.**
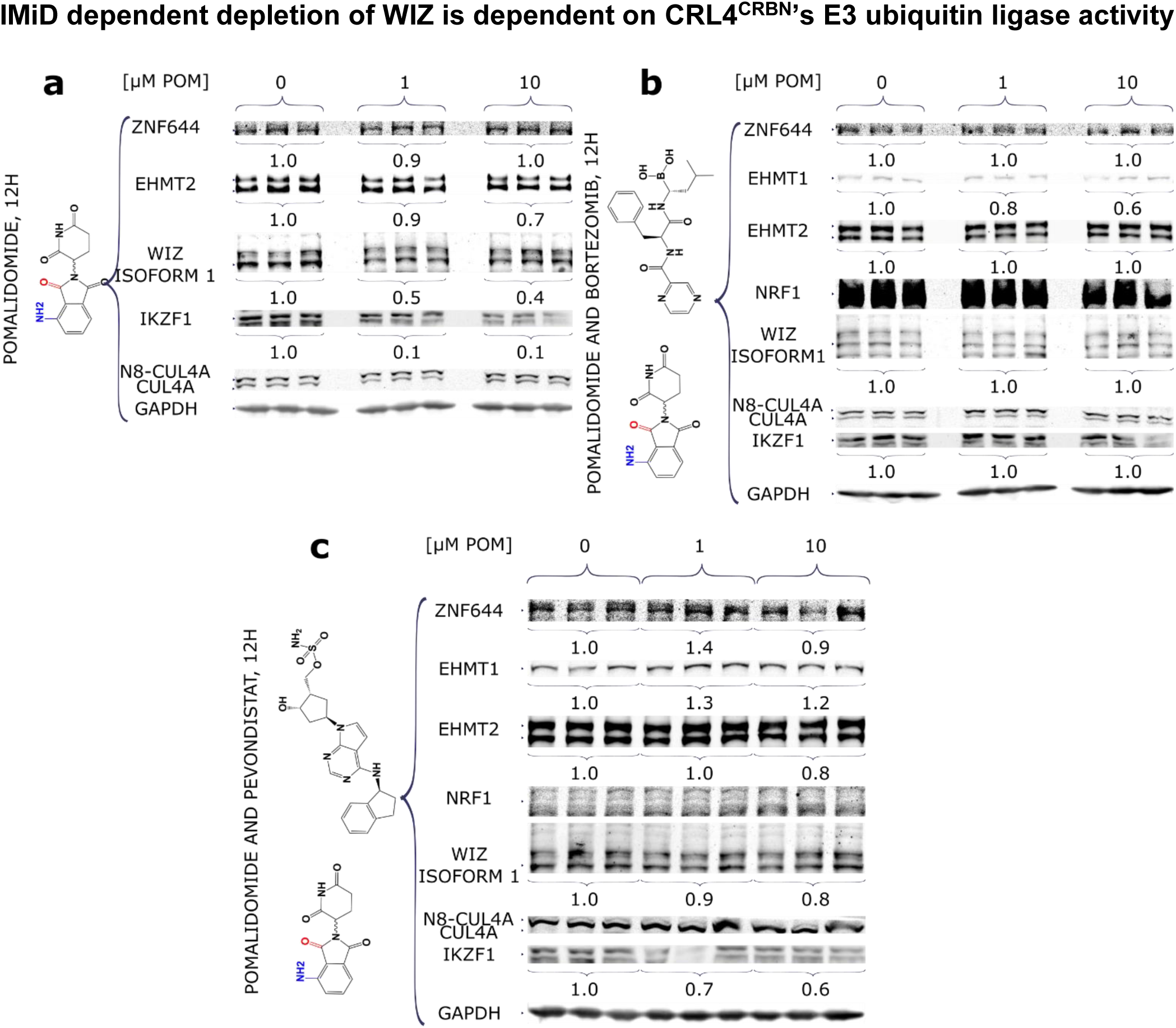
IMiD induced degradation of WIZ is dependent on E3 ubiquitin ligase activity and proteasome degradation. L363 cells were maintained in RPMI-1640 with 10% FBS before treatment with pomalidomide at 0, 1, or 10 µM in biological triplicate. They were cotreated with either (**a**) DMSO, (**b**) the proteasome inhibitor bortezomib, or (**c**) the Nedd8 conjugation inhibitor pevonedistat (38, 39). Cell lysates were separated by SDS-PAGE and immunoblotted against the indicated antibodies. NRF1 was used as a positive control for the proteasome inhibitor (40). IKZF1 was a positive control for IMiD treatment. GAPDH was the loading control. The action of pevonedistat was confirmed by its effect on CUL4 neddylation.

### IMiD dependent depletion of WIZ protein levels was due to enhancing the rate of decay

Although we found IMiD dependent depletion of WIZ required both CRBN and CRL ubiquitin ligase activity (Figs.6b, d and 7a–c), it could be due to increased degradation of WIZ or inhibition of WIZ expression. To decouple these two possible outcomes, we compared the rate of decay by treating cells with or without IMiD in the presence of an inhibitor of translation (Fig.8a-c) (42). We saw that both isoforms of WIZ, as well as IKZF1 experienced a higher rate of decay upon addition of IMiD (Fig.8b-c), consistent with this being due to an increased rate of degradation. We saw that IKZF1 and both isoforms of WIZ, but not EHMT1 and EHMT2, exhibited an increased rate of decay upon addition of IMiD (Fig.8b-c), consistent with this being due to an increased rate of degradation targeting only one subunit.

**Fig. 8.**
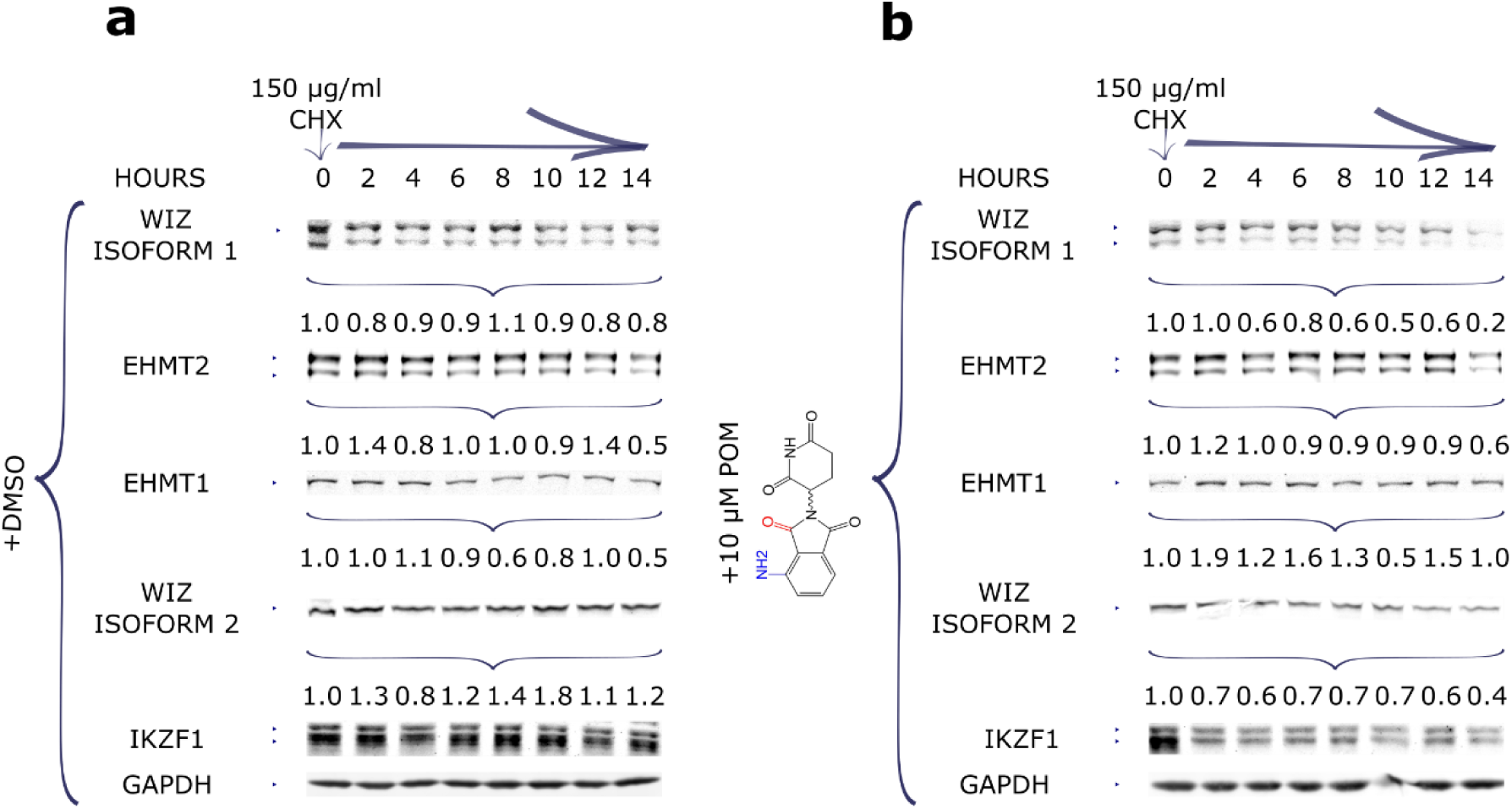
Pomalidomide stimulates the degradation of WIZ, but not other components of the complex. MM.1s cells were maintained in RPMI-1640 before treatment with cycloheximide (150 µg/mL) (**a**) without or (**b**) with 10 µM of pomalidomide. Cells were harvested at 0, 2, 4, 6, 8, 12, and 14 hours, lysed, and separated by SDS-PAGE and immunoblotted for WIZ, IKZF1, EHMT1, EHMT2, and GAPDH.

## Discussion

Targeted protein degradation is rapidly becoming an area of intense interest for the biopharmaceutical industry (43). The overall goal for researchers in this area is to design a drug that selectively targets a particular protein of interest for ubiquitin-dependent degradation via the proteasome. One particular class of drugs, the IMiDs, achieves its effects through selective destruction of specific neosubstrates via the CRL4^CRBN^ ubiquitin ligase pathway (14, 20, 21). Interestingly, IMiDs that differ from each other in subtle ways elicit the degradation of different sets of overlapping neosubstrates. To better understand the biological effects of IMiDs and to gain insight into the mechanisms by which IMiDs recruit substrates to CRBN, we employed an affinity purification-mass spectrometry approach to identify proteins that bound CRBN in an IMiD-dependent manner. This led to the identification of a dozen proteins, including the previously-described CRL4^CRBN^ neosubstrates IKZF1, IKZF3, ZFP91, ZNF692 and CK1*α* (14, 20, 21, 25, 28). In addition, we identified a number of zinc finger proteins, including WIZ. Note that while this work was being prepared for submission, Thomä and colleagues reported the interaction of a large collection of zinc fingers proteins with CRBN, including those reported here (44).

We show here that WIZ is a *bona-fide* neosubstrate for CRL4^CRBN^. In addition to increasing association of WIZ with CRBN, IMiDs also induced depletion of WIZ that was dependent on the CRBN protein, CRL activity (as determined with the pan-CRL inhibitor pevonedistat), and proteasome activity. Importantly, depletion of WIZ was due to an increase in its rate of degradation. The effect of IMiDs on WIZ degradation was selective, in that only IMiDs with an amino group at the 4’ position (lenalidomide and pomalidomide) induced degradation, with pomalidomide showing substantially stronger activity. This is consistent with prior observations that subtle differences in IMiD structure can have powerful discriminating effects on substrates such as CK1*α* (25, 30). In addition, we observed IMiD-dependent depletion of WIZ in L363 but not in THP-1 cells. We do not know the reason for this selectivity. However, it is not due to the inability of WIZ to bind transfected CRBN in an IMiD-dependent manner in these cells (e.g. see Fig.2b) and it is not due to absence of CRL4^CRBN^ activity in THP-1 cells because IKZF1 was efficiently downregulated by IMiDs in these cells. This selectivity points to the potential to develop therapeutic agents that induce degradation of target proteins in a cell type specific manner.

Our data, combined with other recently reported results, raise the question of whether the effects of IMiDs *in vivo* are more complex than originally envisioned, and could potentially be influenced by depletion of multiple proteins in addition to the original neosubstrates IKZF1, IKZF3, and CK1α (14, 20, 21, 25). A great deal of effort is being invested in developing proteolysis-targeting chimeric molecules (PROTACs) that, like IMiDs, induce the degradation of specific proteins to achieve a therapeutic effect (43, 45). A result we observed here that is of particular relevance to those efforts is that IMiD induced depletion is highly selective for a single subunit of a multisubunit complex. WIZ associates with EHMT1, EHMT2, and ZNF644 to form a gene regulatory complex (36, 37), and all four of these proteins were recruited to CRBN in the presence of IMiD. However, only WIZ was degraded. This is consistent with the previously observed subunit selectivity of the ubiquitin proteasome pathway both *in vivo* and in reconstituted systems (46, 47). Harnessing the power of subunit-selective degradation to remodel cellular machines to achieve a highly selective therapeutic outcome holds promise for the development of a suite of next-generation medicines.

### Experimental Procedures

#### A. Materials

Bortezomib (B-1408) was purchased from LC Laboratories. Pomalidomide (P0018-25MG) and *N*-ethyl maleimide (E3876-5G) were purchased from Sigma Aldrich. Lenalidomide (HY-A0003) and pevonedistat (905579-51-3) were purchased from MedChem Express. All were prepared as single-use DMSO stocks and stored at −20^°^ C before usage. Laemmli Buffer (1610737) was purchased from Biorad. BSA (9998S) was purchased from Cell Signaling Technology.

#### B. Antibodies

GAPDH (SC-365062) antibody was from Santa Cruz Biotechnology. CUL4a (2699S), DDB1 (A300-462A), EHMT1 (A301-642A-M), and ZNF644 (A301-642a-M) antibodies were from Bethyl Laboratories. CRBN (HPA045910) and FLAG (F1804) antibodies were from Sigma Aldrich. WIZ (ab92334), ZNF521 (ab156271), CK1*α* (ab108296), and EHMT1 (ab185050) antibodies were from Abcam. PATZ1 (PA5-30478), ZNF684 (PA5-40984), and ZFP91 (PA5-43064) antibodies were from Thermo Fischer Scientific. Anti-Rabbit IgG IR800 (926-32211) was purchased from Li-COR Biosciences. Anti-Mouse IgG IR680 (A10038) was purchased from Invitrogen.

#### C. Cell lines

MM.1s and THP-1 cells were purchased from ATCC. L363 cells were provided by Francesco Parlati, Calithera Biosciences, South San Francisco. Cells were grown in RPMI-1640 (ATCC formulation), with 10% heat-inactivated fetal bovine serum (FBS), 2 mM glutamine, and penicillin-streptomycin. THP-1 cells were supplemented with 50 µM β-mercaptoethanol. Cell lines were tested for mycoplasma using Lonza’s MycoAlert Mycoplasma Detection kit and authenticated by Laragen using PowerPlex 16 system, as well as the Cell Line ID.

#### D. Western blot analysis

MM.1s, THP-1, or L363 cells for each sample, as well as a replicate to measure protein levels were harvested by centrifugation and resuspended in pre-warmed RPMI-1640 at 1 million cells/mL. They were then seeded onto 24 well plates, in 1.5 mL total volume. Pomalidomide was added from a 1000x stock. Cell pellets were collected by centrifugation and flash frozen for processing later. A separate replicate to normalize protein concentration was washed twice with PBS before being flash frozen. Samples were then solubilized in 200 µL of Laemmle buffer supplemented with fresh β-mercaptoethanol and protease inhibitor cocktail. Cell lysates were sonicated (5 s; 10%) and cleared by centrifugation before boiling for 3 minutes. Boiled lysates were cleared by centrifugation (14,000 xg). Relative protein concentrations were calculated based on samples lysed in 2% SDS in the BCA assays, and the total lysate volume was adjusted according to that. Samples were loaded onto 4–12% protein gels. Following electrophoresis, gels were transferred to nitrocellulose for 3 hours at 70 V, with stirring.

#### E. RNAi-mediated knockdown

siRNAs were purchased from Santa Cruz Biotechnology. Cells were seeded at a final concentration of 500,000/mL (THP-1 cells), and transfected with Opti-MEM and Lipofectamine RNAimax, as previously described. Cells were harvested after 48 and 72 hours and analyzed for efficiency of knockdown.

#### F. Cycloheximide Chase

MM.1s cells were seeded at 106 cells/mL in complete RPMI-1640 with 10% FBS in 24 well plates. Cells were treated with 0 or 10 µM pomalidomide, along with 150 µg/mL of cycloheximide to initiate the chase. Cells were harvested at the indicated times and subjected to immunoblot analysis.

#### G. Mass Spectrometric sample preparation

Eluted samples were reduced in 3 mM TCEP for 20 min. Reduced samples were alkylated with 10 mM NEM for 30 min at room temperature in the dark. Reduced and alkylated samples were digested with Lys-C (Wako) at a 1:200 ratio for 4 hours at room temperature. Partially digested lysates spiked with 1 mM CaCl_2_ were diluted with 50 mM Tris (pH 8.0) down to 2 M urea before being digested with sequence grade trypsin (Promega) diluted 1:100, at 37°C overnight in the dark. The reaction was quenched after 15 hours by adding trifluoroacetic acid (TFA) to 0.1%. Lysates were cleared via centrifugation at 4000xg for 15 minutes. Peptides were desalted using a 500 mg capacity Sep-Pak column that was prehydrated with 7 column volumes of acetonitrile (21 mL), followed by an equilibration step with 7 column volumes of Buffer A (0.1% TFA in H_2_O) (21 mL). Cleared peptides were loaded onto the resin by gravity flow, washed with 7 column volumes of Buffer A, followed by 3 column volumes of Wash Buffer (0.1% TFA, 5% acetonitrile in H_2_O). Desalted peptides were eluted with 2 column volumes of Elution Buffer (0.1% TFA, 40% acetonitrile in H_2_O) (6 mL). The resulting peptide sample was frozen by storing at 80°C for at least 1 hour and dried via lyophilization.

#### H. NanoLC-MS/MS analysis

The dried immunoprecipitated peptides were resuspend 0.2% formic acid, 2% acetonitrile, nanoLC grade 97.8% H_2_O and subjected to proteomic analysis using an EASY 1000 nano-UPLC (Thermo Fisher Scientific) connected on-line to an Orbitrap Fusion mass spectrometer with a nanoelectrospray ion source (NanoFlex, Thermo Scientific). Peptides were separated with a 120 min gradient using a 15 cm silica analytical column with a 75 µm inner diameter packed in-house with reversed phase ReproSil-Pur C18AQ 3 µm resin (Dr. Maisch GmbH, Amerbuch-Entringen, Germany). The flow rate was set to 350 nl/min, and the gradient was as follows: 2-6% (7.5 min), 6-25% (82.5 min), 25-40%(30 min), 40-100% (1 min) B (0.2% formic acid, 80% acetonitrile, 19.8% nanoLC grade H_2_O). The mass spectrometer was set to collect data in a data-dependent (top speed) mode, switching automatically between full-scan MS and tandem MS acquisition. Samples were analyzed by Higher Collision Energy (HCD) fragmentation.

#### I. Data Analysis

Raw data was searched using MaxQuant (48, 49), version 1.6.5.0. Precursor mass tolerance was set to 4.5 ppm after recalibration and fragment tolerance was 0.5 Da. Oxidation of methionine and protein N-terminal acetylation were specified as variable modifications and carbamidomethylation of cysteine was specified as a fixed modification. Trypsin was specified as the digestion enzyme, with up to two missed cleavages allowed. Spectra were searched against the UniProt human database (93591 entries) and a contaminant database (246 entries). Score thresholds were established so that peptide and protein false discovery rates were less than 1% as estimated by a target-decoy approach. Match between runs and iBAQ quantitation (50) were enabled. iBAQ parts per million (ppm) abundances were calculated by multiplying individual protein iBAQ abundances by 10^6^ and dividing by the sum of all non-contaminant protein iBAQ values.

## Supporting information

Knockdown

## Acknowledgements

Thanks to Rati Verma, Jennifer Mamrosh, Emily Blythe, Thang Nguyen, and David Sherman for scientific discussion. Thanks to Thang Nguyen for MM.1s CRBN KD and parental cell lines, and preparing samples in Fig. 2. H.H.Y was supported by NSF GRFP fellowship. R.J.D. was an investigator at the Howard Hughes Medical institute, and this work was supported by HHMI and a gift from Amgen.

