## Supplementary material for "Single subunit degradation of WIZ, a lenalidomide- and pomalidomide-dependent substrate of E3 ubiquitin ligase CRL4^CRBN^": Knockdown

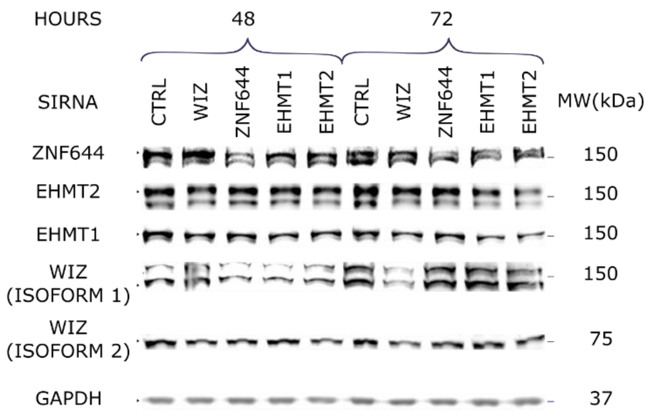


**Fig. S1.** THP-1 cells were transiently transfected with an siRNA against WIZ, EHMT1, EHMT2, or ZNF644. Cells were harvested after 48 or 72 hours and immunoblotted against different commercially available antibodies.
